# Within-patient and global evolutionary dynamics of *Klebsiella pneumoniae* ST17

**DOI:** 10.1101/2022.11.01.514664

**Authors:** Marit A. K. Hetland, Jane Hawkey, Eva Bernhoff, Ragna-Johanne Bakksjø, Håkon Kaspersen, Siren Irene Rettedal, Arnfinn Sundsfjord, Kathryn E. Holt, Iren H. Löhr

**Affiliations:** Department of Medical Microbiology, Stavanger University Hospital, Stavanger, Norway; Department of Biological Sciences, Faculty of Mathematics and Natural Sciences, University of Bergen, Bergen, Norway; Department of Infectious Diseases, Central Clinical School, Monash University, Melbourne, Australia; Research Section Food Safety and Animal Health, Department of Animal Health and Food Safety, Norwegian Veterinary Institute, Oslo, Norway; Department of Paediatrics, Stavanger University Hospital, Stavanger, Norway; Faculty of Health Sciences, University of Stavanger, Stavanger, Norway; Department of Medical Biology, Faculty of Health Sciences, UiT – The Arctic University of Norway, Tromsø, Norway; Norwegian National Advisory Unit on Detection of Antimicrobial Resistance, Department of Microbiology and Infection Control, University Hospital of North Norway, Tromsø, Norway; Department of Infection Biology, Faculty of Infectious and Tropical Diseases, London School of Hygiene & Tropical Medicine, London, United Kingdom; Department of Clinical Science, Faculty of Medicine, University of Bergen, Bergen, Norway

**Keywords:** *Klebsiella pneumoniae*, ST17, *in vivo* evolution, infection, colonisation, global dynamics

## Abstract

*Klebsiella pneumoniae* sequence type (ST) 17 is a global problem clone that causes multidrug-resistant (MDR) hospital infections worldwide. In 2008-2009, an outbreak of MDR ST17 occurred at a neonatal intensive care unit (NICU) in Stavanger, Norway. Fifty-seven children were colonised. We observed intestinal persistence of ST17 in all of the children for up to two years after hospital discharge. Here, we investigated the within-host evolution of ST17 in 45 of those children during long-term colonisation and compared the outbreak with 254 global strains. Ninety-two outbreak-related isolates were whole-genome sequenced. They had capsule locus KL25, O locus O5 and carried yersiniabactin. During within-host colonisation ST17 remained stable with few single nucleotide polymorphisms, no acquisition of antimicrobial resistance (AMR) or virulence determinants, and persistent carriage of a *bla*_CTX-M-15_-encoding IncFII(K) IncFIB(K) plasmid (pKp2177_1). The global collection included ST17 from 1993-2020 from 34 countries, that were from human infection (41.3%), colonisation (39.3%) and respiratory specimens (7.3%), from animals (9.3%), and from the environment (2.7%). We estimate that ST17 emerged mid-to-late 19th century (1859, 95% HPD 1763-1939) and diversified through recombinations of the K and O loci to form several sublineages, with various AMR genes, virulence loci and plasmids. There was limited evidence of persistence of AMR genes in any of these lineages. A globally disseminated sublineage with KL25/O5 accounted for 52.7% of the genomes. It included a monophyletic subclade that emerged in the mid-1980s, which comprised the Stavanger NICU outbreak and 10 genomes from three other countries, which all carried pKp2177_1. The plasmid was also observed in a KL155/OL101 subclade from the 2000s. Three clonal expansions of ST17 were identified, all were healthcare-associated and carried either yersiniabactin and/or pKp2177_1. To conclude, ST17 is globally disseminated and associated with opportunistic hospital-acquired infections. It contributes to the burden of global MDR infections, but many diverse lineages persist without acquired AMR. We hypothesise that non-human sources and human colonisation may play a crucial role for severe infections in vulnerable patients, such as preterm neonates.

**Impact statement:** *Klebsiella pneumoniae* is an opportunistic pathogen that frequently causes hospital-associated multidrug-resistant (MDR) infections. Infections with *K. pneumoniae* strains that are resistant to third generation cephalosporins and/or carbapenems are considered a major public health threat, as there are limited treatment options available. Some MDR *K. pneumoniae* clones, including ST307 and ST258, are global problem clones because they disproportionately contribute to the burden of MDR infections and are common causes of such infections and/or outbreaks in hospitals around the world. Here we describe another such clone, ST17, which caused an MDR outbreak in our neonatal intensive care unit, affecting 57 children. We found that this clone underwent minor within-host evolution during two years of long-term gastrointestinal colonisation after hospital discharge. We then investigated the evolutionary history of ST17 globally, as it had not previously been studied. When we compared ST17 isolates from around the world with the isolates from our hospital, we discovered that whilst ST17 with antimicrobial resistance has contributed to outbreaks in other neonatal care units, it also frequently causes infections or colonises humans and non-human sources without any antimicrobial resistance.

**Data summary:** The genome sequences generated in this study have been deposited at the European Nucleotide Archive under BioProject PRJEB36392. The BioSample accession numbers and associated metadata for the genomes are available in Table S1. The completed annotated genome assembly of Kp2177 is available in GenBank under accession numbers CP075591 (chromosome), CP075592 (pKp2177_1), CP075593 (pKp2177_2) and CP075594 (pKp2177_3). The global dataset of 300 ST17 genomes is available for interactive viewing in Microreact at https://microreact.org/project/kpst17.

## Introduction

The Gram-negative bacteria *Klebsiella pneumoniae* are a common cause of infections in both healthcare and community settings (1, 2). Notably, they are a major pathogen responsible for bacterial outbreaks in neonatal intensive care units (NICUs) (3–5). Several sequence types (STs) of *K. pneumoniae* are associated with multidrug-resistance (MDR) (6). Some clones carry genes encoding extended-spectrum beta-lactamases (ESBLs) or carbapenemases and co-linked resistance determinants, resulting in severely limited treatment options (5). Some MDR clones remain localised, e.g. within a healthcare or community setting, whilst others have spread across continents and become global problem clones (5). One such clone is *K. pneumoniae* ST17, which has been detected in several countries, often with an MDR phenotype causing difficult-to-treat hospital infections (5, 7–9). ST17 has also been found colonising healthy humans in the community setting (10–12), in animals (13, 14) and in the environment (15–17).

From November 2008 to April 2009 an outbreak caused by ESBL-producing (*bla*_CTX-M-15_) ST17 occurred in a NICU at Stavanger University Hospital, Norway (8). The outbreak was first discovered when ESBL-producing *K. pneumoniae* was recovered from three neonates: one clinical sample from conjunctiva and two routine screening samples. Subsequent screening revealed that 92% (22/24) of the patients in the ward were colonised with ESBL-producing *K. pneumoniae*. The index isolate of the outbreak (Kp2177) was isolated from breast milk from one of the mothers. During the outbreak, a total of 57 children were colonised with ST17. Only one of them developed a bloodstream infection (BSI), which was successfully treated with meropenem. A follow-up study revealed intestinal persistence of the ST17 strain in all 57 children after hospital discharge, all carrying a *bla*_CTX-M-15_-encoding IncFII(K) IncFIB(K) plasmid, for up to two years (median 12 months) (18, 19).

Here, we first characterised the genomic diversity and *in vivo* evolution of ST17 within and between 45 NICU patients in Stavanger, to study the evolution of this strain during long-term gut colonisation. As the *bla*_CTX-M-15_ ST17 strain persisted for up to two years in these children, we wanted to examine how the *in vivo* evolution compared to the evolutionary dynamics of ST17 on a global scale. Whilst ST17 has been observed in several countries and settings, specific analyses to determine its global dynamics have not been reported previously. We therefore carried out a literature and genome search to assess the global presence of ST17, followed by core genome evolutionary analyses of 300 ST17 strains from around the world, that were isolated from human infections and colonisation, animals, plants and water.

## Methods

### Collection and sequencing of the Stavanger NICU outbreak isolates

During the outbreak in the Stavanger NICU, 57 children were colonised. They were examined for intestinal persistence over a two-year period (19), and for 45 of the children, we had two ST17 isolates available for genomic comparison in this study. For each child, one sample was isolated during the NICU stay (between January and March 2009) and another after hospital discharge (between June 2009 and November 2010). The duration of colonisation between the two isolates from each child ranged 3-21 months (median 11). All of the colonising isolates were from faecal swabs except one nasal swab isolate. We performed whole-genome sequencing using Illumina MiSeq (n=46) or HiSeq (n=46) technology to produce 300 bp or 150 bp paired-end short-read sequences of in total 92 isolates: the 45 NICU and 45 follow-up isolates, the one BSI isolate and the index (breast milk) isolate Kp2177. BioSample accessions are given in Table S1. To generate a reference genome for the outbreak, Kp2177 was also sequenced with the Oxford Nanopore Technologies (ONT) platform, and hybrid assembly was performed to resolve the complete genome sequence (deposited in GenBank under accessions CP075591-94; details in Supplementary Methods).

### Global dataset of ST17 genomes

A literature search querying PubMed for titles or abstracts containing the word “ST17” with and without “*Klebsiella pneumoniae”* resulted in 36 peer-reviewed articles that contributed novel data on *K. pneumoniae* ST17 (as of November 2021), but only 29 ST17 isolates were whole-genome sequenced and available for download. We therefore also included ST17 genomes from other sources: a recent study analysed 13,156 publicly available *K. pneumoniae* genomes and identified their STs (20). Those that were confirmed ST17 and had <500 assembled contigs, known year of collection and sample source (n=149) were included in our global dataset. Additionally, we included 64 ST17 genomes from the recent SpARK study (17) and 41 ST17 genomes from Norwegian surveillance studies that were not linked to the Stavanger NICU outbreak (11, 21–23). For analyses of global dynamics, we also included 46 of the Stavanger NICU outbreak genomes, resulting in a global dataset of 300 genomes (BioSample accessions are given in Table S2).

### Genome assembly and genotyping

All short-read Illumina sequences were filtered with TrimGalore v0.6.6 (https://github.com/FelixKrueger/TrimGalore) to exclude low-quality reads and adapter-contamination before *de novo* assembly was performed with Unicycler v0.4.8 (24), using SPAdes v3.13.0 (25) and Pilon v1.23 (26), with default settings. Detection of antimicrobial resistance (AMR) and virulence genes was based on analysis of genome assemblies using Kleborate v2.2.0 (20), which uses a manually curated version of CARD v3.1.13 for acquired AMR determinants. We also used SRST2 v0.2.0 (27) to query the unassembled short-reads directly, using the same database. If a gene was missing in the Kleborate analysis of assemblies but found with SRST2 analysis of reads, we considered it present in the genome. Capsule (K) and lipopolysaccharide (LPS) O-antigen serotype predictions were performed with Kaptive v2.0.0; as low-confidence calls are most likely due to assembly-quality, we reported the best matching K and O loci regardless of confidence call, except for those called with low-confidence as KL107 which were designated unknown (28). Plasmid replicons were detected using Abricate v1.0.1 (https://github.com/tseemann/abricate) with the PlasmidFinder database v2021-01-13 (https://bitbucket.org/genomicepidemiology/plasmidfinder_db).

### Between and within-host comparison of the Stavanger NICU outbreak genomes

To identify single nucleotide polymorphisms (SNPs) between and within the outbreak genomes, core genome alignments were produced with RedDog v1beta.11 (https://github.com/katholt/RedDog) by aligning the raw reads of the 92 outbreak genomes against Kp2177. Within-host SNPs were defined as SNPs between the NICU and follow-up genome of a child, from the overall core genome alignment. Bowtie2 v2.3.5.1 (29) was used for sequence alignment and SAMTools v1.9 (30) to identify SNPs. Further details are given in the Supplementary Methods. RAxML v8.2.12 (31) was used to infer a core genome maximum likelihood (ML) phylogeny from the chromosomal alignment of 113 SNPs. To estimate a substitution rate (SNPs/year) for the outbreak, we divided the number of SNPs against Kp2177 from each genome by the difference in isolation time.

To identify genes that had been gained or lost between the NICU and follow-up genomes, we characterised their pangenome with Panaroo v1.2.7 (32), using annotations from Prokka v1.14.6 (33) as input. As all the genomes belonged to the same outbreak strain and we wanted to identify any changes in gene content, we ran Panaroo using sensitive mode with the protein family identity set to 90%.

### Phylogenetic inference and molecular dating of the global ST17 collection

We performed ML and Bayesian phylogenetic analyses of 300 genomes to compare the Stavanger NICU outbreak (n=44 follow-up faecal, 1 blood, 1 breast milk) with ST17 from other geographical locations and collection years (n=254), and to estimate the evolutionary rate of the clone. We generated a core chromosomal SNP alignment of the genomes against Kp2177 with RedDog as described in the Supplementary Methods. Gubbins v3.1.6 (34) was used to identify and filter recombinant SNPs from the alignment, which was then passed to RaxML v8.2.12 (31) to infer an ML phylogeny. We investigated the relationship between the root-to-tip distance in the ML tree and the years of isolation using linear regression in TempEST v1.5.3 (35). To estimate the evolutionary rate and dated phylogeny, we performed Bayesian phylodynamic analysis with BEAST2 v2.6.5 (36) on a subset of 145 of the 300 genomes, which were selected to capture the diversity across the clades in the full ML tree as detailed in the Supplementary Methods. The temporal signal of the clone was confirmed by performing date-randomisation tests (Figure S1).

### Statistical analyses

All statistical analyses were performed with R version 4.0.2 (2020-06-22) (37). Paired data were analysed with a two-tailed Wilcoxon paired signed rank test. Comparisons of groups were analysed with Wilcoxon signed rank test and binomial data were compared with Chi square test. P-values <0.05 were considered statistically significant.

## Results

### The Stavanger NICU outbreak

The index isolate (Kp2177) of the Stavanger NICU outbreak, which was isolated from breast milk, belonged to ST17, had capsule locus KL25 and O locus O5, and consisted of a 5.38 Mbp chromosome and three plasmids (pKp2177_1, 182 Kbp, IncFII(K) and IncFIB(K); pKp2177_2, 83 Kbp, IncR and IncFIA(H1); and pKp2177_3, 3 Kbp, no plasmid replicon marker detected). A total of 5,303 genes were annotated in the genome. pKp2177_1 harboured the ESBL gene *bla*_CTX-M-15_, the beta-lactamase gene *bla*_TEM1-D_ and the aminoglycoside resistance gene *aac(3)-Iia*. pKp2177_2 carried *strAB, dfrA14, sul2*, and *cat2* encoding resistance towards streptomycin, trimethoprim, sulphonamides, and phenicols, respectively. Of known virulence factors, yersiniabactin (*ybt* 16) was chromosomally encoded on an ICE*Kp12* mobile genetic element. No other siderophores or hypermucoviscosity-encoding genes were present.

All of the colonising NICU (n=45) and subsequent follow-up isolates (n=45) belonged to the same ST17 strain, with KL25/O5 and ≤8 SNPs compared to the index Kp2177 isolate. The three Kp2177 plasmids were conserved in all 45 NICU genomes, with high sequence coverage (98.3%-100%, Table S1). The AMR and virulence genes in Kp2177 were also present in all of the NICU genomes, except for one that did not harbour *cat2* (however the follow-up genome for this child did). There was more variation in the follow-up genomes, which were collected up to 21 months (median 11) after initial colonisation (Figure 1). The *bla*_CTX-M-15_-encoding plasmid pKp2177_1 was persistent in all of the strains with minimal variation (≤1 SNP and 68.8%–100% sequence coverage, median 100%). The pKp2177_2 and pKp2177_3 plasmids varied in coverage and AMR-gene content and were missing in 16% (n=7) and 11% (n=5) of the 45 follow-up genomes. In six of the follow-up genomes we detected plasmid replicon markers that were not present in Kp2177 or in the NICU genomes. However, none of the 90 colonising genomes encoded additional AMR or virulence determinants not present in Kp2177. Some children were given antibiotics during their NICU stay or after hospital discharge (Table S3). No significant difference in the number of AMR genes was observed between the children that had received antibiotics (n=34) and those that did not (n=11) (p=0.92).

**Figure 1.**
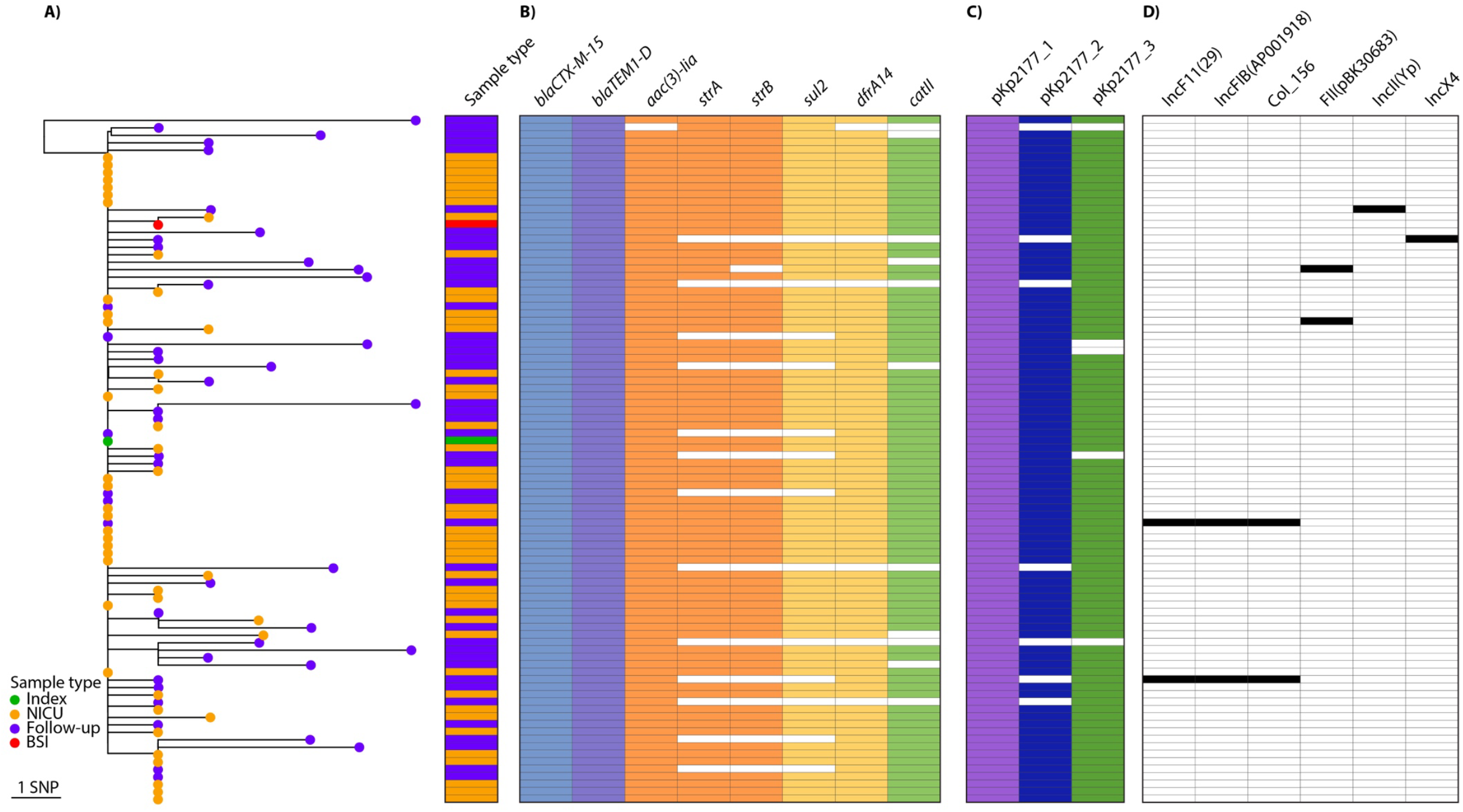
Phylogeny of 92 *Klebsiella pneumoniae* ST17 outbreak isolates and their gene content. **A)** Midpoint-rooted maximum likelihood phylogeny of the 45 NICU (orange tip) and 45 follow-up (purple) isolates from colonisation and the BSI isolate (red) against the outbreak index isolate Kp2177 (green). **B)** Presence (colour) or absence (white) of AMR genes as listed in the columns (blocks are coloured by drug class). **C)** Presence (colour) or absence of the Kp2177 plasmids and **D)** of acquired plasmid replicon markers.

Only one child developed a systemic infection (KN_0144A) (8). The BSI genome also belonged to the Kp2177 strain and was highly similar (1 SNP, *wza*-G178A), with no acquisition of genes that were unique to this genome. The follow-up genome for this patient (KN_0144A-L), which was isolated from gut colonisation 14 months after the BSI occurred, had the most SNPs of any in the outbreak (n=10). None of them were present in other outbreak genomes (Figure 1A and Table S4), nor in the global ST17 dataset (see below).

### Diversity and evolution within and between hosts

To assess the diversity and *in vivo* evolution of the outbreak strain over a 21-month period, we performed a core genome phylogenetic analysis using Kp2177 as the reference genome. In total, 120 unique SNPs were identified, 113 of which were in positions in the chromosome, 3 in pKp2177_1 and 4 in pKp2177_2 (full list of SNPs and their consequences are given in Table S4). The 45 NICU genomes differed from Kp2177 by 0-3 SNPs (median 1). In contrast, the 45 follow-up genomes differed from the index by 0-8 SNPs (median 2) and showed significantly more evolutionary divergence from Kp2177 (p<0.0001). Based on the chromosomal alignment and the time between Kp2177 and the isolate collection dates, we calculated a substitution rate for the outbreak of 2.60 SNPs/year (95% CI 1.96–3.24), or 4.84×10^−7^ SNPs/year/site (95% CI 3.64×10^−7^–6.02×10^−7^).

We observed few genome changes within-hosts during up to 21 months of colonisation. Within-host SNPs were detected in 84.4% (38/45) NICU and follow-up pairs (Table S3). The number of SNPs in each child ranged from 0-10 (median 2), of which up to 6 were nonsynonymous SNPs (nsSNPs) and the rest synonymous or intergenic. Most of the SNPs (91.7%, n=110/120) occurred only once, in one genome each. However, in eight genes, >1 SNP was observed, in different genomes (Table S4). Notably, two genes had several SNPs: *barA* and *uvrY*, that make up the BarA-UvrY two-component system (TCS) (38). In *uvrY*, there were two SNPs leading to different amino acid changes in the same codon (G76W, G76R). In *barA*, four nsSNPs were present, two resulting in premature stop codons (Q260*, R305H, V311G and Q466*). These six mutations were variably present in nine genomes from children that were colonised >6 months (median 1.2 years). Of these mutations, only *uvrY*-G76R was also present in the global ST17 dataset (see results section below), in one genome (KP_NORM_URN_2013_95870), but an additional 12 nsSNPs in these two genes were present across the global phylogeny in 19 genomes.

Among the 91 colonising genomes there were 5,742 unique genes. Of those, 76.5% (n=4,393) were core, i.e. present in all of the genomes. The accessory genes were either very common (60.3% of the 1,349 accessory genes were present in >90% of the genomes) or rare (32.2% were present in <5% of genomes). Overall, there was a significant within-host increase in the number of genes in the follow-up (median 5,279; range 5,081-5,428) compared to the NICU genomes (median 5,253; 5,175-5,306) (p=0.005, Figure S2). Within each child, between 6 and 311 (median 71) gene gain or loss events occurred during the colonisation period. In five of the 45 children, more than 150 genes had been gained during colonisation; none of them encoded known AMR or virulence determinants or plasmid replicon markers. There were also two children in which over 150 genes had been lost.

### ST17 is globally disseminated and highly diverse

To understand the ST17 variation in the Norwegian children in the context of the global diversity of this strain, we included 300 ST17 genomes for comparative and phylogenetic analyses. The genomes were globally distributed, including isolates from 34 countries in 12 United Nations defined geographical regions over a 28-year period (1993–2020) (Figure 2 and Table S2). The majority were from humans (88%, n=264/300): infection (n=124, most from blood [n=79] or urine [n=43]), colonisation (n=118, most from faeces [n=109, the Stavanger NICU outbreak accounted for 44 of these]), and respiratory sites with unknown infection status (n=22). The genomes from non-human sources (12%, n=36/300) came from animals (n=26, from dogs, broilers, a chicken, cows, a fly, pigs and turkeys), marine bivalves (n=2), different sources of water (n=7) and a sugar cane (n=1). The animal genomes were sampled from intestinal isolates, except one from a chicken meat isolate.

**Figure 2.**
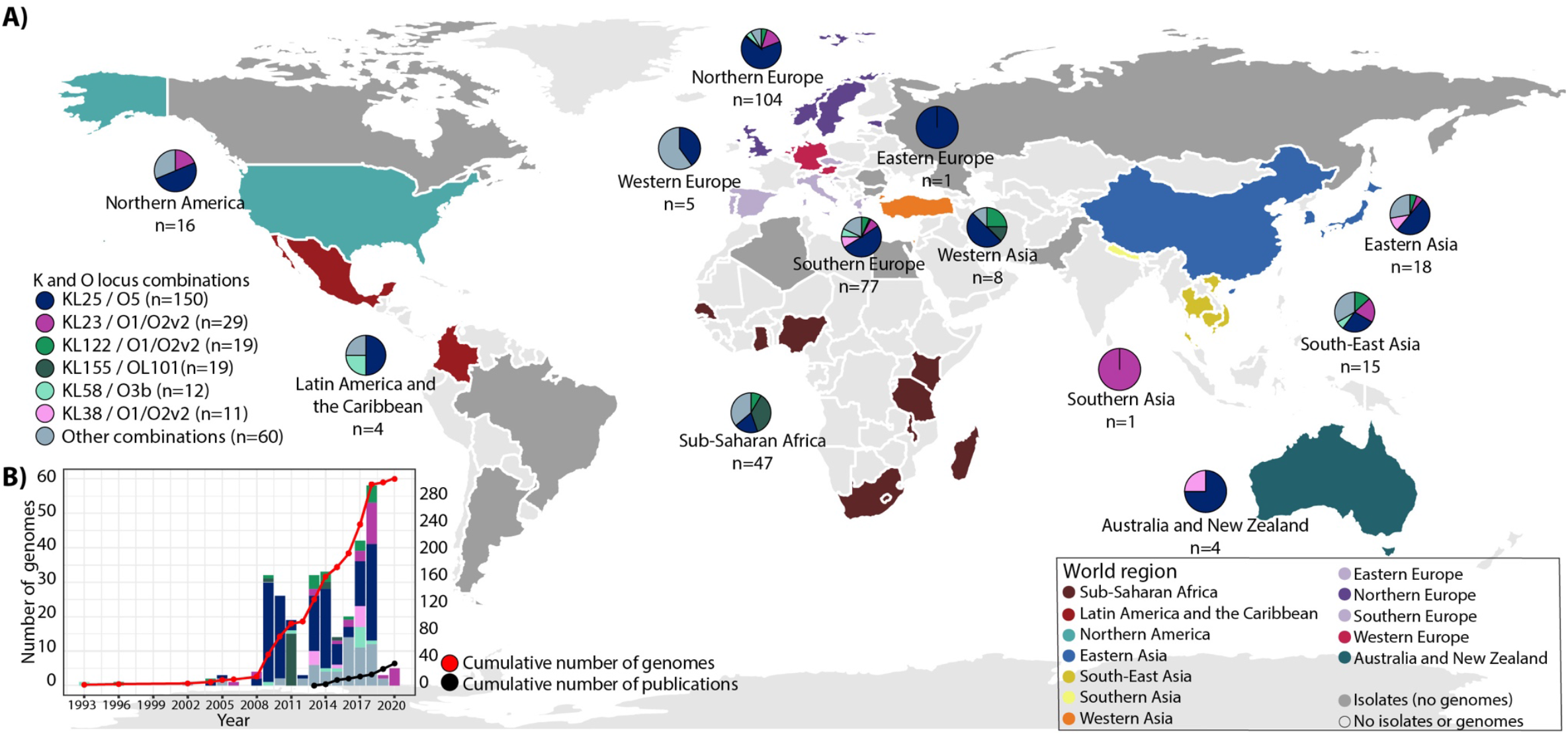
Geographical distribution of *Klebsiella pneumoniae* ST17 genomes, their K and O loci, and reports of ST17. **A)** World map highlighting countries of collection for ST17 genomes that were available for download (coloured by world region as in Figure 3) and countries with isolates reported but not included in the global dataset (coloured dark grey). The pie charts show the distribution of K and O loci per world region (n=number of genomes), highlighting loci with ≥10 genomes. **B)** Number of ST17 genomes by collection year (bars, coloured by KL/OL) and cumulative number (red line), and cumulative number of reports of ST17 in PubMed abstracts or titles (black line) as of November 2021.

A maximum likelihood phylogeny of the 300 genomes was created (Figure 3, also available for interactive viewing at https://microreact.org/project/kpst17). Due to the size and diversity of the dataset, 145 representative sequences were selected to estimate the evolutionary rate of ST17 using Bayesian modelling in BEAST2 (Figure S3) (see details in Supplementary Methods). The Bayesian analysis of 11,345 SNPs in 145 genomes indicated that ST17 emerged in 1859 (95% HPD 1763-1939) with an estimated evolutionary rate of 4.04× 10^−7^ substitutions/site/year (95% HPD 2.30× 10^−7^–5.79× 10^−7^), equivalent to 2.17 SNPs per year. Since its emergence, the clone has undergone several recombination events (the ratio of nucleotide substitutions introduced by recombination vs point mutation was estimated as r/m=3.27), which were centred in a hotspot surrounding the K and O loci (Figure 4A). These recombinations have led to a diversification of ST17 strains by creating distinct sublineages with different combinations of K and O loci. In total, there were 8 O and 27 K loci (including unknown KLs), and 34 combinations of these, among the 300 ST17 genomes (Figure 4B). Over half (52.7%, n=158/300) of the genomes belonged to a clade that had KL25/O5 (except for two genomes with KL2/O2v1, one KL166/O5 and five unknown KL25/O5) that emerged in 1943 (95% HPD 1905-1972). This clade was found in 19 countries across 11 world regions (all except Southern Asia), and in all sample types. The genomes in the remaining sublineages of ST17 were found in 29 countries (all world regions except Eastern Europe) and had a much larger variety of K and O loci (32 combinations), with KL23/O1/O2v2 (n=29), KL155/OL101 (n=19) and KL122/O1/O2v2 (n=18) being the most prevalent (Figure 2). KL23/O1/O2v2 and KL122/O1/O2v2 were intermingled with genomes carrying other combinations of K/O loci, whilst KL155/OL101 formed a monophyletic clade.

**Figure 3.**
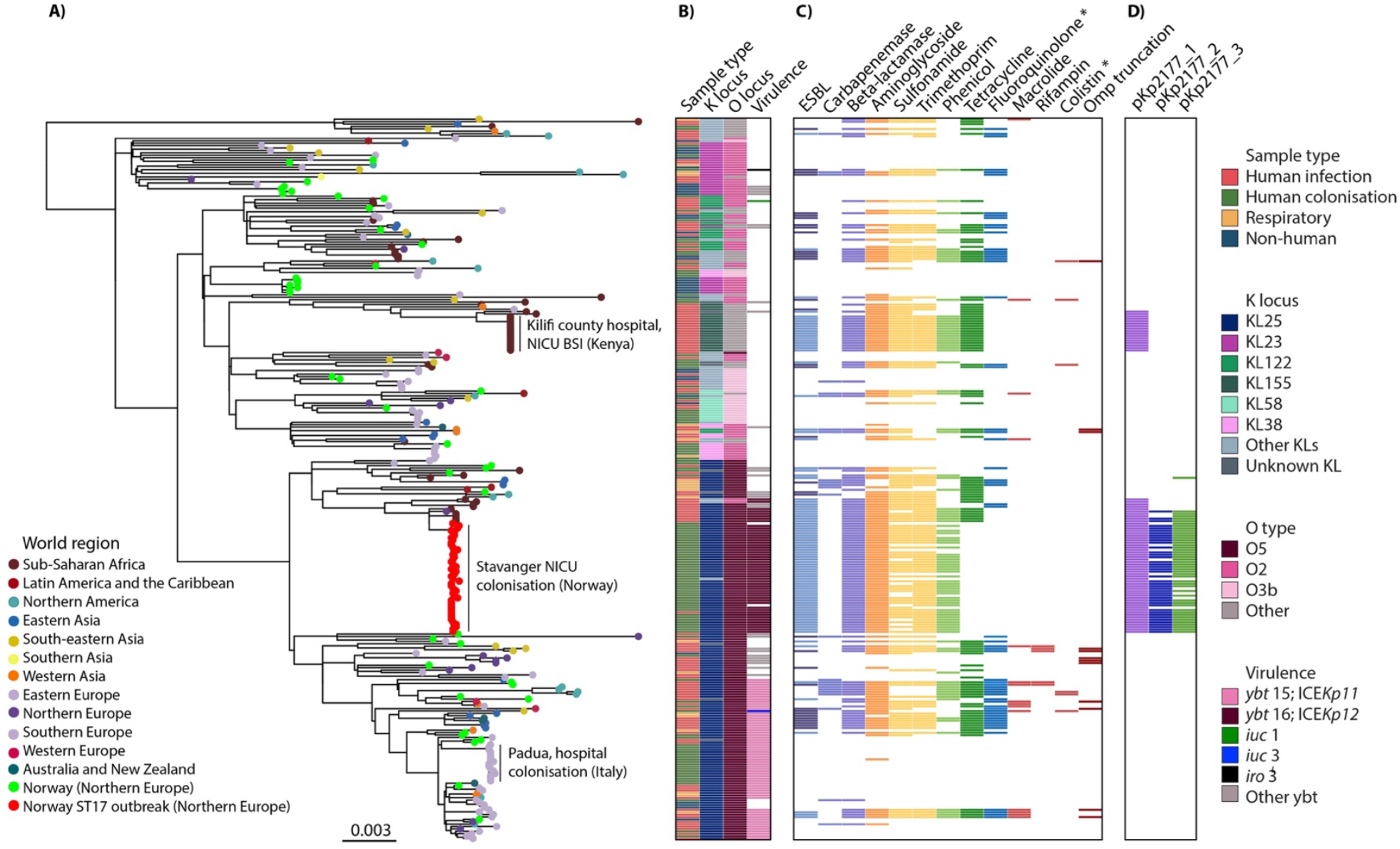
Global phylogeny of 300 *Klebsiella pneumoniae* ST17 genomes. **A)** Maximum likelihood tree with tips coloured by world region of collection. Additionally, the Stavanger NICU outbreak genomes are coloured red and other genomes from Norway green. The three clonal expansions that were observed in ST17 are labelled. **B)** Sample type and loci as indicated in the column names. The most prevalent loci are indicated in the inset legend. **C)** Presence (colour) or absence (white) of genes encoding resistance to the listed antimicrobial resistance (AMR) drug classes (blocks are coloured by drug class). Lighter colour in the ESBL column indicates *bla*_CTX-M-15_. **D)** Presence (colour) or absence (white) of the Kp2177 plasmids. Several plasmid replicon markers were present across ST17, these are shown in Figure S4. More details about the metadata and genotypes are available for interactive viewing at https://microreact.org/project/kpst17. * Acquired genes and mutations.

**Figure 4.**
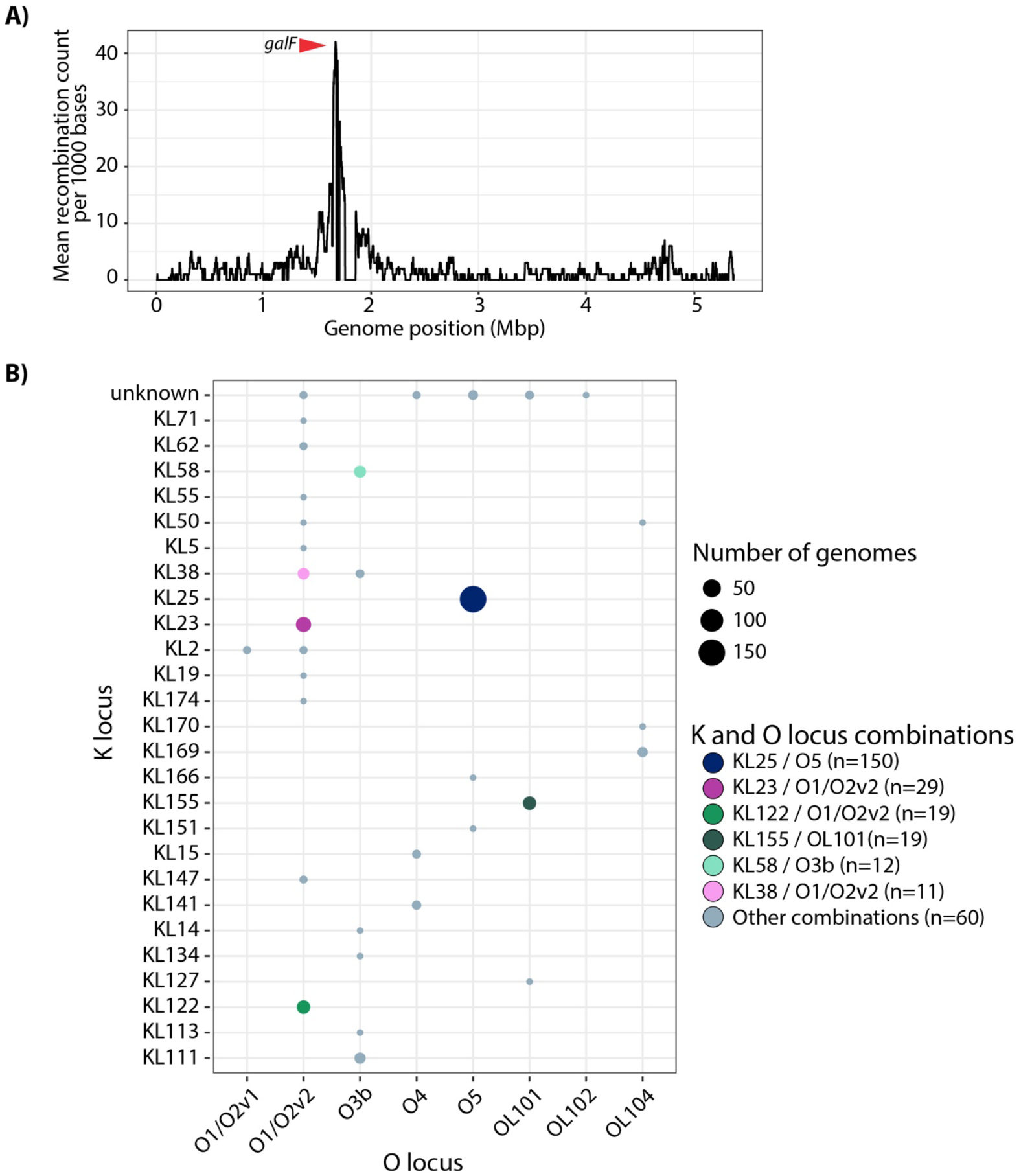
Recombinations and K/O loci diversity. **A)** Recombination counts per base calculated over non-overlapping windows of 1000 base pairs. There is a peak in the recombination count, which surrounds the K and O loci. The red arrow indicates the gene *galF*, which is the 5’ K locus gene. **B)** Combinations of K and O loci found among the 300 genomes. Bubbles indicate the number of genomes and are coloured by the K/O combinations as in Figures 2 and 3.

The dominant KL25/O5 clade was also distinct from the rest of the tree by high prevalence of yersiniabactin: 84.2% (133/158) of the genomes in this clade carried a yersiniabactin locus, compared to 7.0% (10/142) of the genomes in the rest of the tree. The majority of the yersiniabactin loci were *ybt* 15, located on ICE*Kp11*, (n=63, complete or truncated) and *ybt* 16, on ICE*Kp12*, (n=52, complete or truncated), which formed distinct subclades in the KL25/O5 sublineage (see Figure 3). Other yersiniabactin loci (n=10) were sporadically present across the tree, in 28 genomes, including in these subclades. There were few hypervirulence-associated genes, but aerobactin (*iuc 1* and *3*) or salmochelin (*iro 3*) with incomplete hypermucoviscosity *rmp* genes were present in three genomes.

Several different AMR determinants (n=81) and plasmid replicon markers (n=44) were detected in the ST17 genomes (Table S2 and Figure S4). In the overall collection, the number of AMR genes ranged from 0 to 19 (median 3), with 42.7% (128/300) genomes having no AMR genes present at all and 48.3% (145/300) being MDR (carried resistance genes to ≥1 antimicrobial agent in >3 antimicrobial classes). We observed no differences in the number of AMR determinants or MDR genomes between the samples from humans (infection, respiratory and colonisation). However, significantly fewer AMR genes were detected in the non-human isolate genomes (non-human: median 0, range 0-11 vs human: median 4.5, range 0-19, p<0.0001). Of the AMR-encoding genomes, 127 carried ESBL genes; *bla*_CTX-M-15_ (n=92), *bla*_CTX-M-14_ (n=20), *bla*_CTX-M-27_ (n=1), *bla*_CTX-M-63_ (n=1), *bla*_SHV-12_ (n=13), *bla*_SHV-2_ (n=3), *bla*_SHV-5_ (n=2), *bla*_SHV-7_ (n=1), *bla*_SHV-24_ (n=1), and 25 carried carbapenemase-encoding genes; *bla*_KPC-2_ (n=10), *bla*_NDM-1_ (n=6), *bla*_KPC-3_ (n=4), *bla*_OXA-48_ (n=3), *bla*_NDM-7_ (n=1), *bla*_OXA-181_ (n=1) and *bla*_VIM-1_ (n=1). The AMR genes were found across many sublineages, but there was limited evidence of AMR-persistence within subclades.

### Clonal expansions of ST17 were hospital-associated and carried yersiniabactin or a blaCTX-M-15-encoding plasmid

Only three clonal expansions (≥15 genomes) were observed across the global dataset, and all were hospital-associated (see Figure 3). One of these included the Stavanger NICU outbreak, which formed a monophyletic clade. Its closest neighbours were a cluster of four identical NICU blood isolates from Kilifi, Kenya (2009; 47 SNPs from Kp2177), five blood isolates from Nigeria (2009; 40 SNPs, 2015-2016: 143-831 SNPs) and one from the United Kingdom (UK) (2011; 38 SNPs) (Figure S5). The most recent common ancestor (MRCA) for this cluster was 1984 (95% HPD 1967-1998). These genomes carried the *bla*_CTX-M-15_-encoding pKp2177_1 plasmid that was persistent in the Stavanger NICU outbreak (≤1 SNP and 100% sequence coverage), and were highly similar, except for four genomes from Nigeria; two that had KL2/O2v1 instead of KL25/O5, and two that carried *ybt* 13 (on ICE*Kp2*) rather than *ybt* 16 (on ICE*Kp12*).

The *bla*_CTX-M-15_ pKp2177_1 plasmid was present (≤2 SNPs and ≥95% sequence coverage, median 100%) in one other clonal expansion of ST17, in a cluster of 17 genomes, which all had KL155/OL101, of which only one genome encoded yersiniabactin (*ybt 9* on ICE*Kp3*) (see Figure 3). These were invasive isolates from Kilifi, Kenya in 2011 (n=15, 1 SNP between them), Nigeria in 2015 (n=1, 89 SNPs from the ones from Kenya) and Ghana (n=1, 90 SNPs) in 2015, with an MRCA of 2000 (95% HPD 1990-2007).

The last of the three clonal expansions consisted of 16 isolates from gut colonisation in a hospital in Padua, Italy during 2018 (17) (see Figure 3). These had KL25/O5 and carried *ybt* 15 (on ICE*Kp11*) but no AMR genes (except one with *aac(6’)-Ib-Suzhou*). In 2017-2018, 48 other ST17s were isolated from different sources in Padua, Italy. These were not related to the clonal expansion in the local hospital, but were found across the entire phylogeny, carrying a range of KL/OLs, and with varying presence of yersiniabactin and AMR genes. The same was the case for the Norwegian ST17 genomes that were not part of the Stavanger NICU outbreak (n=43).

To see if the pKp2177_1 plasmid had spread in Norway between STs, we screened 3,212 non-ST17 *K. pneumoniae* genomes for the presence of the *bla*_CTX-M-15_ pKp2177_1 plasmid (see Supplementary Methods). These covered 1,076 STs from multiple human and non-human sources in Norway between 2001 and 2020. We found only one genome that carried this plasmid, with 100% sequence coverage; a clinical ST307 urine isolate from 2013, previously described in (39).

## Discussion

In this study we investigated the global dynamics of *K. pneumoniae* ST17 and the *in vivo* evolution within and between children that were colonised following an outbreak in an NICU in Stavanger, Norway in 2008–2009. During up to 21 months of colonisation, few changes (0-10 SNPs) occurred in the core genome, and the majority were present in only one genome each. However, there was one SNP that was shared among several follow-up genomes and it was also found in one genome in the global dataset. This was in the gene *uvrY* (G76R). UvrY is a response regulator that is part of a TCS with BarA, a sensor kinase, that has been described in uropathogenic *Escherichia coli* (38). Across the outbreak and global ST17, several independent nsSNPs were seen in these two genes. Pernestig et al. (38) found that mutations in either of those genes can affect survival in long-term competition cultures; when using media with gluconeogenic carbon sources, *E. coli* with knockout mutations in *barA* or *uvrY* had a clear growth advantage over the wild type, whereas when using media with carbon sources feeding into the glycolysis, the wild type had the growth advantage. Another study found that homologues of these genes in other bacteria led to decreased virulence in infection models (40, 41). It is possible that the mutations in these genes in ST17, if rendering the genes non-functional, gave an advantage to long-term colonisation in the gut, which is a source of gluconeogenic carbon.

The Stavanger NICU outbreak strain persistently carried a *bla*_CTX-M-15_-encoding IncFII(K) IncFIB(K) plasmid (pKp2177_1) for up to two years, but the presence of other plasmids varied. Whilst some genes were gained during the follow-up period, no additional known virulence or antibiotic resistance encoding determinants were acquired. This could be a result of the children acquiring a more diverse microbiome once they left the hospital, enabling horizontal gene transfer to or from the ST17 strains. The outbreak and the global ST17 had overlapping evolutionary rates, but unlike the relatively stable genomes within the Stavanger NICU outbreak, ST17 exists globally with many sublineages, harbouring various AMR determinants and plasmid replicons, frequently carrying yersiniabactin and in a few cases also the hypervirulence-associated aerobactin and salmochelin loci.

The first detected ST17 was isolated from a pig brain in the UK in 1993 (14), and the earliest ESBL-producing isolate was a *bla*_CTX-M-15_-carrying clinical isolate from Canada in 2002 (42). However, the Bayesian phylogeny analysis suggested that ST17 emerged much earlier than this, in the mid-to-late 19th century. ST17 is older, and has a slower evolutionary rate (4.04× 10^−7^ substitutions/site/year), compared to other global problem MDR lineages that have been estimated, such as ST307 (1.18 × 10^−6^ substitutions/site/year, MRCA 1994), ST258 (1.03×10^−6^, 1995) and ST147-KL64 (1.03×10^−6^, 1994) (39, 43, 44). Since its emergence, recombination has driven diversification of the K and O biosynthesis loci in ST17, which has led to the formation of several sublineages. This is similar to that seen in other MDR clones, including ST258 and ST147 (45).

The MDR clones ST258, ST307 and ST147 frequently encode ESBL or carbapenemase genes (5). Unlike these, ST17 was not strongly associated with any specific AMR genes. In fact, less than half of the ST17 genomes carried resistance to third generation cephalosporins and/or carbapenems. On the other hand, 48.3% (145/300) of ST17 genomes were MDR, carrying various AMR genes, including carbapenemases in 25 genomes and ESBL in 127, of which 92 carried *bla*_CTX-M-15_. MDR strains were found in many sublineages, but there was limited evidence of persistence of any AMR genes within these, nor within specific countries or collection years, with the possible exception of the *bla*_CTX-M-15_-encoding pKp2177_1 plasmid, which was present in two clonal expansions.

One of the sublineages, KL25/O5, which emerged in the mid-1940s, contained over half the ST17 genomes and was globally disseminated. The Stavanger NICU outbreak was located on this sublineage, in a cluster with 10 genomes from Nigeria, Kenya and the UK. They all carried the *bla*_CTX-M-15_-harbouring pKp2177_1 plasmid, with an estimated MRCA in 1984. The plasmid was also seen in another sublineage, KL155/OL101, with 17 genomes from Kenya, Ghana and Nigeria (MRCA 2000). The isolates from the two sublineages in Kenya were from 2009 and 2011, and both were from the same NICU in Kilifi County Hospital (46). In that hospital, ST17 with/without ESBL phenotypes were observed in blood culture isolates almost every year since it first emerged (46), indicating undetected transmission of this strain in the hospital or region, and that the pKp2177_1 plasmid is circulating between ST17 strains. Given the MRCAs of these pKp2177_1 plasmid-encoding clusters and the larger diversity of the pKp2177_1-encoding genomes in Nigeria, it is probable that the plasmid was acquired by an ancestor in Nigeria in the mid-1980s, where it subsequently diversified and transmitted to the other countries, and that the plasmid later transferred into or between these two ST17 subclades, either in Nigeria or Kenya. In a previous study, only minor fitness costs were associated with carriage of pKp2177_1 in the Stavanger NICU outbreak, indicating that the plasmid was well adapted to its bacterial host (18). Further, even though the plasmid carried a complete conjugation machinery, we observed no evidence of plasmid transfer to *E. coli in vitro* (18) or to other bacterial hosts *in vivo* during the two years of colonisation (19). Additionally, only one of 3,212 non-ST17 genomes from Norway carried pKp2177_1, further supporting that pKp2177_1 is well adapted to ST17.

We included two large collections of ST17 from Norway (n=89) and Padua, Italy (n=69) (17). The genomes were spread across the ST17 sublineages and were found in several host species and sample types. This shows that the diversified descendants of the ancestral ST17 co-circulate both within and between ecological niches, even in highly localised geographic regions. Further, it suggests that there have been multiple independent importations of ST17 to these locations, and that the same patterns would be observed for other countries given larger sample sizes, which was also the case for ST307 (39). The NICU outbreaks in the collection indicate that this highly flexible and widely disseminated clone can rapidly colonise neonates in hospital settings. This may be associated with the presence of the *bla*_CTX-M-15_ pKp2177_1 plasmid and/or yersiniabactin, although more data would be needed to confirm this: The 2009 NICU outbreaks in Stavanger, Norway and Kilifi, Kenya carried the pKp2177_1 plasmid and *ybt* 16 on ICE*Kp12*; the 2011 NICU outbreak in Kilifi carried the plasmid but no yersiniabactin locus; and in Padua, Italy, the genomes did not carry the plasmid, but had *ybt* 15 on ICE*Kp11*. In addition to these large hospital transmissions, some small local clusters (≤7 genomes) occurred. These were mainly reported from studies investigating the movement of strains in close communities, e.g. circulation of ST17 between animals (22) or between animals and people in the community (13), where the same strain was present in animals and humans, indicating that animals may be reservoirs for ST17.

It has been estimated that around half of healthcare-associated infections caused by *K. pneumoniae* arise from opportunistic colonising strains infecting immunocompromised and vulnerable patients, such as neonates (1, 47, 48). Consistent with this, the ST17 NICU outbreak in Stavanger mainly caused intestinal colonisation and only one systemic infection occurred, after colonisation. In Kilifi, Kenya, only invasive infections with ST17 were reported, however colonisation was not screened for and it is likely that it existed without being detected. The ability of ST17 to cause infection may be related to opportunity, such as host factors and environment, rather than its own intrinsic properties. ST17 isolates have predominantly been reported in studies aiming to characterise Gram-negative bacteria or *K. pneumoniae* isolated from given populations (often ESBL- or carbapenemase-producing), as minor STs (i.e. incidental findings), and not because they were causing disease outbreaks (Table S5). For example, a study of AMR Gram-negative bacteria in low- and middle-income countries found that *K. pneumoniae* was the main cause of neonatal sepsis (49). In that study, ST17 accounted for 3.1% (8/258) of the *K. pneumoniae* isolates. Some outbreak reports do exist, and notably they mainly concern NICU outbreaks, e.g. in Norway (8), Kenya (46) and China (7, 9). Taken together, this indicates that ST17 strains typically colonise a host before opportunistically causing infections, rather than causing clonal expansions of infectious outbreaks within wards, although there are exceptions in highly vulnerable host populations such as those found in NICUs.

In conclusion, ST17 is a globally disseminated clone that frequently colonises both humans and non-human sources, and opportunistically causes infections, often in neonates. During within-host colonisation for up to 21 months, ST17 in the Stavanger NICU outbreak remained comparatively stable, with few SNPs and no acquisition of AMR or virulence determinants. Globally, ST17 is highly diverse and harbours a range of AMR genes, virulence loci and plasmids. It contributes to the global burden of MDR infections but is also frequently found without AMR. The ESBL-encoding pKp2177_1 plasmid or yersiniabactin may offer some advantage in clonal expansions of ST17 strains in hospitals, but more data would be needed to assess this. Given the movement of the pKp2177_1 plasmid, the presence of similar ST17 strains in animals and humans, and the deep-branching global phylogeny, we hypothesise that non-human sources and human colonisation play a crucial role for severe infections in vulnerable patients, such as in preterm neonates.

## Author contributions

MAKH, JH, KEH, AS and IHL conceptualised the study. SR collected samples and data from the Stavanger NICU outbreak. EB and RJB performed whole-genome sequencing. HK contributed bacterial genomes from animals. MAKH performed the genomic analyses. MAKH wrote the first draft of the manuscript. All authors contributed to data interpretation, reviewed and edited the manuscript, and have read and agreed to the published version of the manuscript.

## Acknowledgements

We wish to thank Olav Natås and Knut Øymar for their support during the NICU outbreak and follow-up study, and Umaer Naseer for his contribution and expertice related to previous experimental plasmid studies. We thank Edward Feil for providing then unpublished genomes from the SpARK One Health consortium project, and others who have made genomes and metadata from their studies publicly available in online repositories. We also wish to thank Benjamin Silvester, Zoe Dyson, Kelly Wyres, Margaret Lam and Ryan Wick for valuable discussions. This work was presented at the 13^th^ International Meeting on Microbial Epidemiological Markers (IMMEM XIII) in Bath, UK in September 2022.

