## Supplementary methods and figures for "Within-patient and global evolutionary dynamics of *Klebsiella pneumoniae* ST17"

### Contents

**Supplementary Methods:**
See pages 1-5 of this document for methods supplementary to the main text.

**Supplementary Figures:**
**Figure S1:** Temporal signal amongst ST17 genomes.
**Figure S2:** Paired gene gain/loss events between NICU and follow-up genomes.
**Figure S3:** Dated phylogeny of 145 *Klebsiella pneumoniae* ST17 genomes from around the world.
**Figure S4:** Global phylogeny of 300 *Klebsiella pneumoniae* ST17 genomes and plasmid content.
**Figure S5:** Zoom-in of the clade with the Stavanger NICU outbreak genomes and closely related genomes that all carried the $bla_{CTX\text{-}M\text{-}15}$ pKp2177_1 plasmid on the KL25/O5 sublineage.

**Supplementary Tables:**
Please see separate Excel file for the Supplementary Tables:
**Table S1:** Stavanger NICU outbreak genome information, BioSample accessions and genotypes.
**Table S2:** Global ST17 dataset with BioSample accessions, metadata and genotypes.
**Table S3:** Within-host colonisation time, number and type of SNPs, and number of gene gain-loss events during the Stavanger NICU outbreak.
**Table S4:** Stavanger NICU outbreak SNPs, SNP type and affected gene products.
**Table S5:** Summary of literature that mentions *Klebsiella pneumoniae* ST17 in PubMed abstracts or titles, as of November 2021.

### Supplementary methods

#### Collection and sequencing of the Stavanger NICU outbreak isolates

Kp2177, a *Klebsiella pneumoniae* ST17 isolated in November 2008 from the breast milk of a neonate's mother, was previously identified as the likely index isolate for the outbreak. To resolve the sequence of this genome, we whole-genome sequenced the isolate using the same DNA extraction on both Illumina MiSeq and Oxford Nanopore Technologies (ONT) MinION platforms. The ONT sequencing was performed on an r9.4.1 flow cell, basecalled with guppy v3.0.3+ (available for ONT customers at https://nanoporetech.com) and subsampled using FiltLong v0.2.0 (https://github.com/rrwick/Filtlong) by discarding reads shorter than 1 Kbp and removing the 10% worst read bases. The short- and long-read sequences were then hybrid assembled using Unicycler v0.4.8 (1) and subsequently annotated with RAST v2.0 (2).

#### Between and within-host comparison of the Stavanger NICU outbreak genomes

Core genome single nucleotide polymorphism (SNP) alignments were produced with RedDog v1beta.11 (https://github.com/katholt/RedDog) to identify SNPs between and within the outbreak samples. Bowtie2 v2.3.5.1 (3) was used with option --sensitive-local for sequence alignment and SAMTools v1.9 (4) was used to identify SNPs. Filters were applied to exclude allele calls from variant sites with ambiguous base calls, Phred quality <30, read depth <5, or evidence of strand bias. For all variant sites, base calls were extracted from the genomes in the alignment, and core genome positions were identified as positions where ≥95% of the genomes had Phred quality ≥20. Genomes were excluded from the alignments if they had <50% sequence coverage and <5X sequence depth to the reference.

Further, to remove any artefacts caused by sequencing, mapping or assembly errors in the Kp2177 reference genome, we aligned the short-reads of Kp2177 against the hybrid-assembled closed genome of itself, and excluded any SNP calls in positions where the two disagreed (this comprised 12 positions in plasmid pKp2177_2). For the remaining 91 short-read sequenced isolates, we mapped the reads from each isolate against the *de novo* assembly of itself to identify any variable positions (there were none). There were 145 SNPs called against plasmid pKp2177_p1. Of those, 142 were present in only two genomes and they were concentrated in two genes. Upon inspection these were most likely homologs and

were therefore excluded from the final list of SNPs. The final alignments consisted of 113 (chromosome), 3 (pKp2177_1), 4 (pKp2177_2) and 0 (pKp2177_3) SNPs.
Gubbins v3.1.6 (5) was used to identify recombination events in the core chromosomal alignment of variable and invariable sites; there were none. RAxML v8.2.12 (6) was then used to infer a core genome maximum likelihood (ML) phylogeny of the SNPs, using a rapid bootstrap analysis searching for the best-scoring ML tree, a GTR substitution model and GAMMA distribution of rate heterogeneity. This was performed in five independent replicates, and the tree with the highest likelihood was retained (Figure 1).

##### Phylogenetic inference and molecular dating of the global ST17 collection

To assess the global dynamics of *K. pneumoniae* ST17, we included 254 genomes from around the world that were downloaded from GenBank or the European Nucleotide Archive. Fourty-one of these genomes were available only as *de novo* assemblies. To allow them to be included in the phylogenetic analysis, SAMtools wgsim was used to simulate 100 bp paired-end reads without error from the assemblies.

A core chromosomal SNP alignment of 46 Stavanger NICU outbreak genomes (one genomes from each patient; 44 follow-up faecal, 1 blood, 1 breast milk) and the 254 global genomes against the Kp2177 chromosome was generated with RedDog. Five SNPs that were artefacts between the Kp2177 short-reads and the reference genome were excluded. Recombinant regions were filtered from the alignment using Gubbins v3.1.6 (5) with the weighted robinson foulds convergence method. The recombination-free alignment (n=10,313 SNPs) was then passed to RaxML v8.2.12 (6) to infer a core genome ML phylogeny, using a rapid bootstrap analysis searching for the best-scoring ML tree, a GTR substitution model and GAMMA distribution of rate heterogeneity. This was performed in five independent replicates. The final ML scores were highly similar and the phylogeny with the best score was imported to TempEST v1.5.3 (7) to investigate the relationship between the root-to-tip distances in the ML tree and the years of isolation. The slope of the root-to-tip regression under heuristic residual mean squared was negative (-0.0002), indicating low or no molecular clock levels (7). As the dataset was quite large, compared to other dated clones such as ST307 (n=95) and CG147 (n=218) (8, 9), we decided to subset the ST17 global dataset for the temporal analysis and repeated the steps above. We used patristic distances to group the 300 genomes into 125 groups, and picked one isolate from each group to represent the diversity in the tree. Four of the resulting groups had >15 genomes. For these, we additionally picked ≥3 genomes to cover each country within the group. The root-to-tip regression slope was now positive, indicating that a molecular clock signal was present.

BEAST2 v2.6.5 (10) was used for the Bayesian phylogenetic analysis of the final recombination-free alignment of 11,345 SNPs in 145 genomes to estimate the phylogenetic tree and evolutionary rate. We specified exact date of sampling for the genomes where this information was available, and for the genomes that we only knew the year or the year and month of sampling (n=78), we used tip calibration in BEAUTi v2.6.5 (10) to sample the tip dates. We ran four different model combinations through BEAST2, using two clock models (strict and relaxed log normal) and two demographic models (constant population and exponential population). Each model was run in three replicates, to 350 million states until all parameters had >200 ESS and removing 10% of the states was sufficient to remove the burn-in. The model with relaxed log normal clock and constant population size was determined the best fit for the dataset: The rate coefficient of variance in the relaxed clock models did not touch 0, meaning the rate was not the same across the tree, so the relaxed model allowing for variation in rate was selected. The growth rate distribution in the constant exponential model went through 0, suggesting that the population was not growing, therefore the constant coalescent was selected. The trees from this model were subsampled with LogCombiner to 22,500 trees, which were passed to TreeAnnotator to summarise them into a single target maximum clade credibility tree (Figure 3). To confirm the temporal signal of the clone, ten independent date-randomisation tests were performed to show that the estimated rate of the true data did not overlap with those from the randomised dates. There was no overlap between the 95% HPD evolutionary rate of the true and randomised data (Figure S1). To confirm that the prior was not driving the results, the analysis was also performed with sampling from priors only (i.e. without the sequence alignment); there was no overlap with the true data.

#### Comparing pKp2177_1 with non-Norwegian ST17 and non-ST17 Norwegian genomes

To determine the presence of the *bla*$_{CTX-M-15}$-encoding pKp2177_1 plasmid, which was persistently present in the Stavanger NICU outbreak, we used RedDog as described above to perform read mapping against two datasets: The collection of 254 *K. pneumoniae* ST17 genomes not from the outbreak, and a collection of 3,212 *K. pneumoniae* genomes of non-ST17 sequence types from Norway that were isolated between 2001 and 2020. They were from human clinical infection (n=2,073; 868 published in (11) and a further 1,205 not yet published), human gut colonisation of healthy adults (n=481) (12), marine bivalves and seawater (n=97) (13) and animals, including turkey and broilers (n=203) (14) and a further 206 isolates not yet published from turkey, broilers, pigs and dogs. Genomes were considered positive for a plasmid if they had <10 SNPs and ≥80% mapping coverage of the reference, as in (15).

#### Supplementary figures

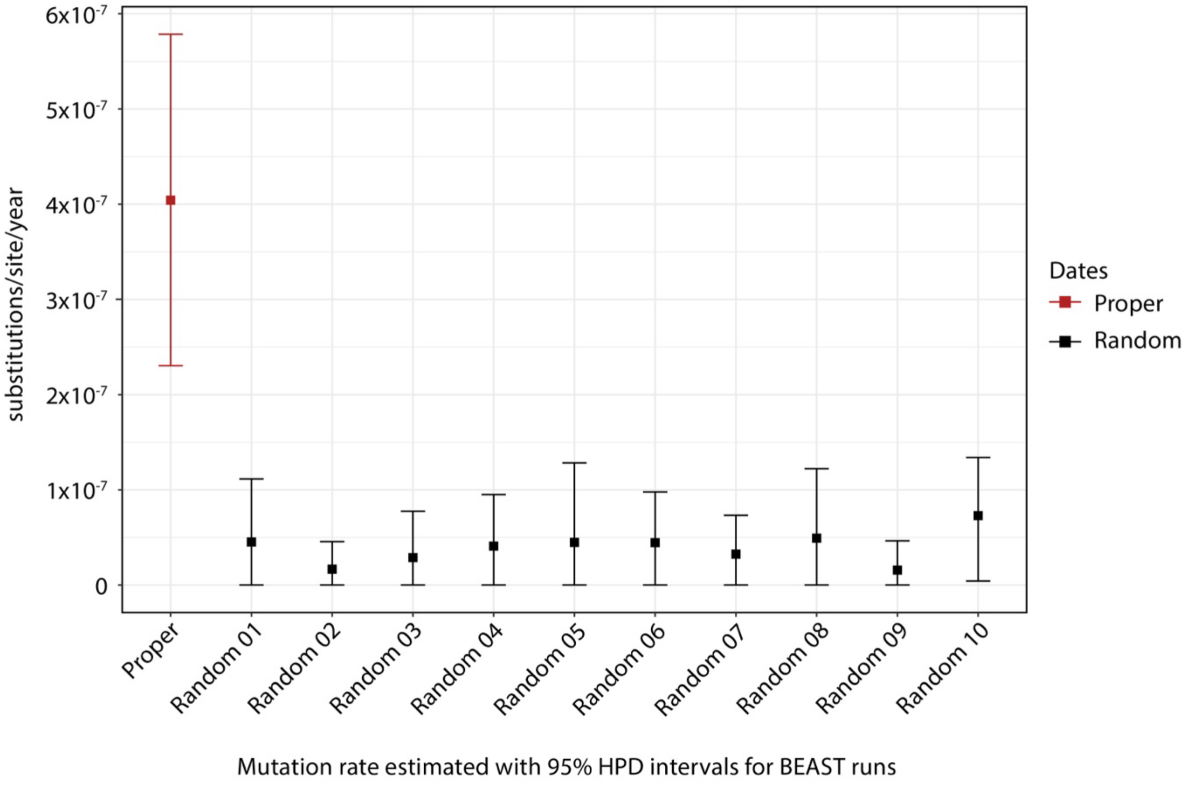


**Figure S1. Temporal signal amongst ST17 genomes.** Posterior distributions for evolutionary rate estimates generated in BEAST2 with the true isolate collection dates (red) and randomised dates (n=10, black).

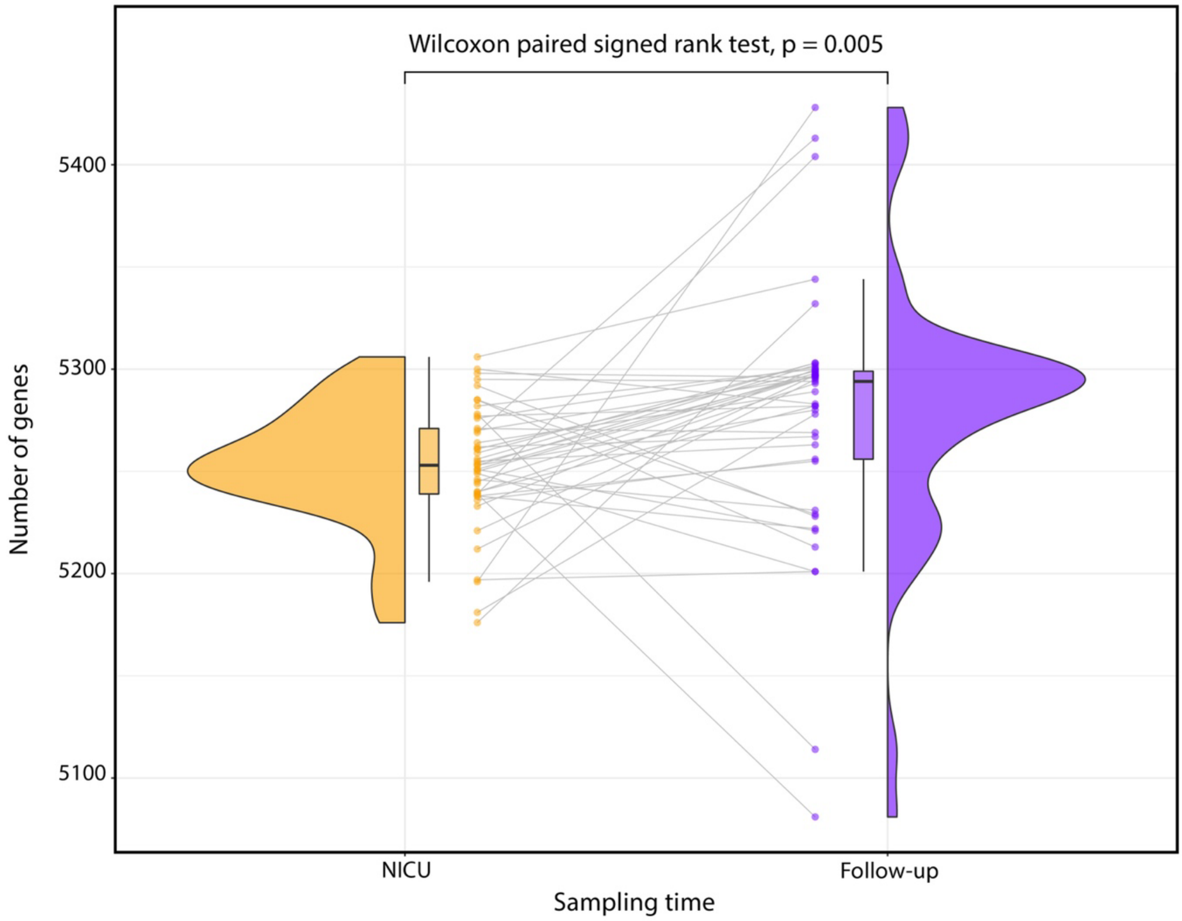


**Figure S2. Paired gene gain/loss events between NICU and follow-up genomes.** Differences in the number of genes between the neonatal intensive care unit (NICU, on the left) and follow-up (taken 3-21 months after colonisation in the NICU, median 11, on the right) isolates. From outer to inner: Density plot, box plot with line indicating median, paired child data points. There was a significant within-host increase in the number of genes in the follow-up genomes than in the NICU ones (p=0.005).

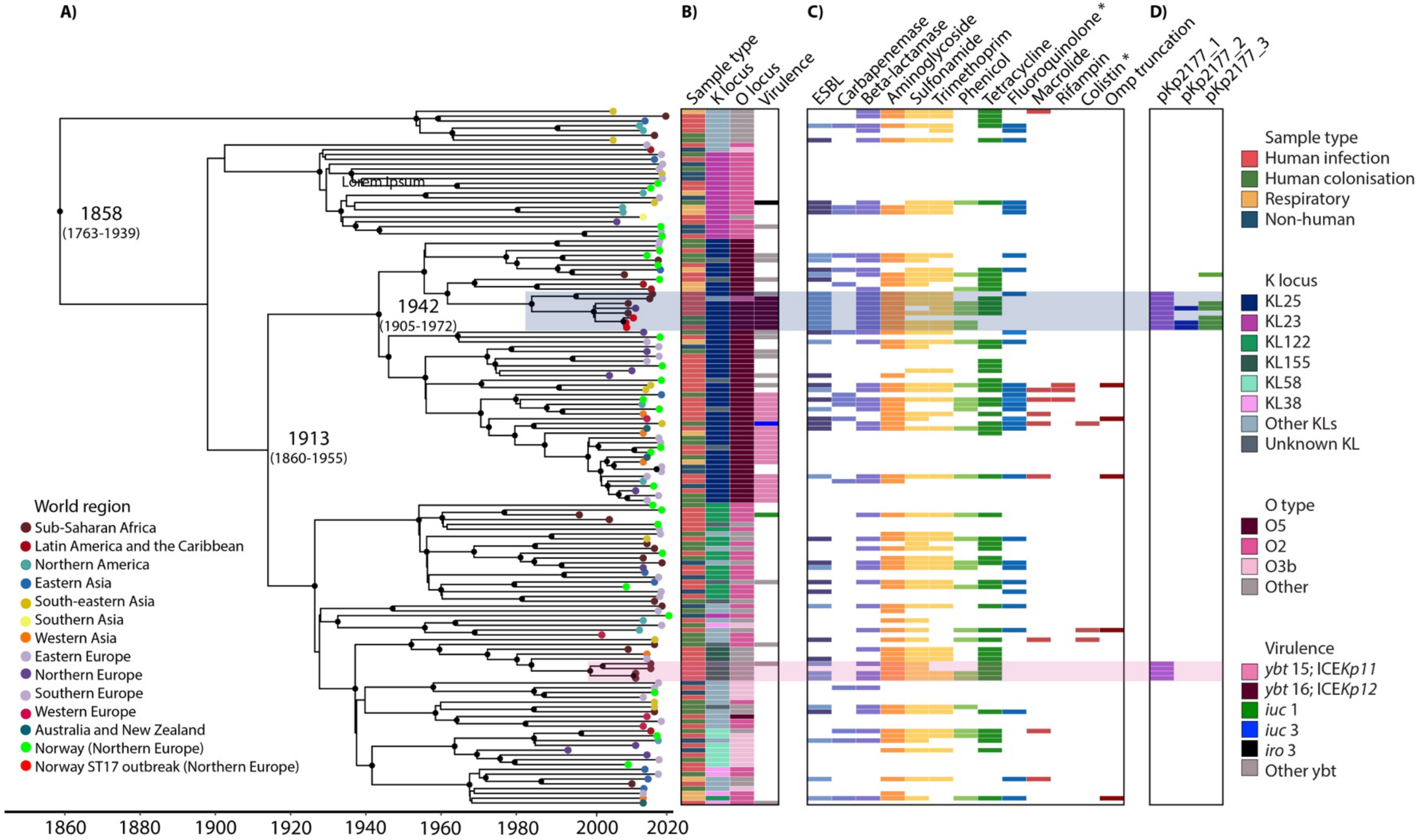


**Figure S3. Dated phylogeny of 145 *Klebsiella pneumoniae* ST17 genomes from around the world. A)** Dated tree with tips coloured by world region of collection. Additionally, the Stavanger outbreak genomes are coloured red and other genomes from Norway green. Black dots on internal nodes indicate ≥95% posterior probability. **B)** Sample type and loci as indicated in the column names. The most prevalent loci are indicated in the inset legend. **C)** Presence (colour) or absence (white) of genes encoding resistance to the listed antimicrobial resistance (AMR) drug classes (blocks are coloured by drug class). Lighter colour in the ESBL column indicates *bla*$_{\text{CTX-M-15}}$. **D)** Presence (black) or absence of the Kp2177 plasmids. The highlighted clades include the representative sequences of the pKp2177_1-harboring clusters: blue = KL25/O5 clade; pink = KL155/OL101. * Acquired genes and mutations.

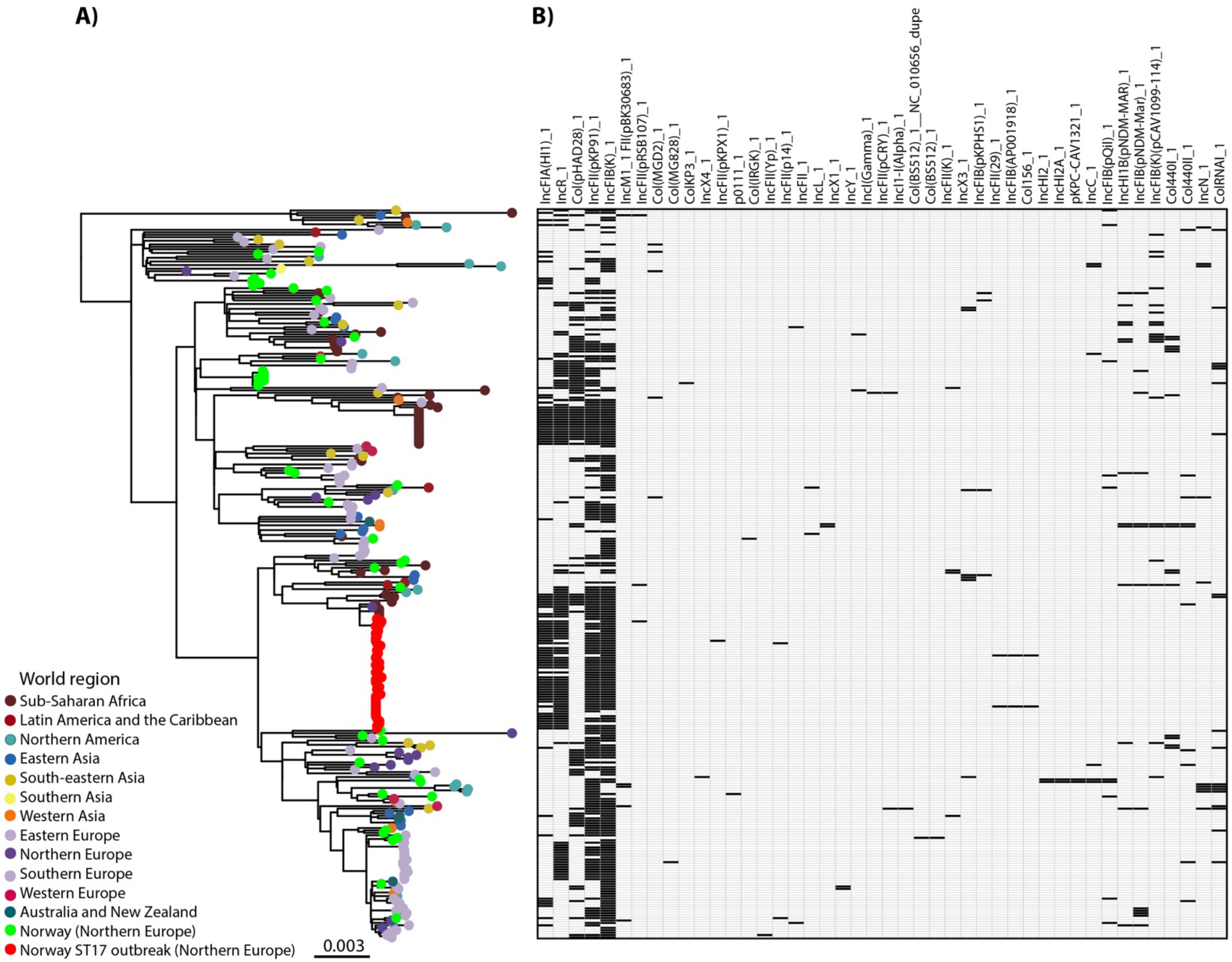


**Figure S4. Global phylogeny of 300 *Klebsiella pneumoniae* ST17 genomes and their plasmid content. A)** Maximum Likelihood tree reproduced from Figure 3. The tips are coloured by world region of collection. Additionally, the Stavanger outbreak genomes are coloured red and other genomes from Norway green. **B)** Presence (black) or absence (white) of all detected plasmid replicon markers in the dataset.

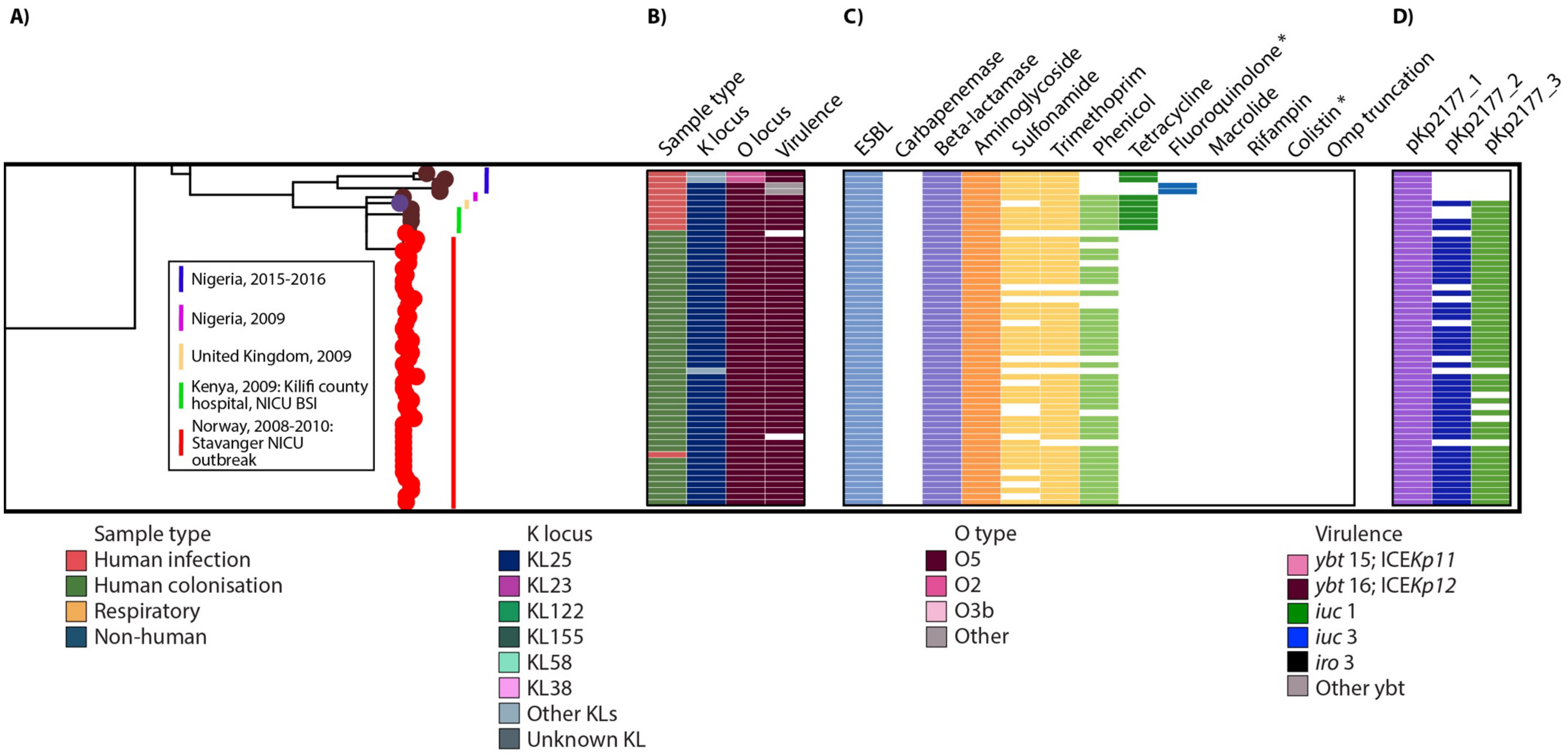


**Figure S5. Zoom-in of the clade with the Stavanger NICU outbreak genomes and closely related genomes that all carried the *bla*$_{CTX\text{-}M\text{-}15}$ pKp2177_1 plasmid on the KL25/O5 sublineage.** The figure is a zoom-in of the maximum likelihood tree in Figure 3. The lines next to the tree tips indicate which country and year the genomes were collected from in the inset legend.
